# Generative single-cell transcriptomics via large language models

**DOI:** 10.64898/2026.01.12.699186

**Authors:** Hongyoon Choi, Haenara Shin, Dongjoo Lee, Daeseung Lee

## Abstract

Single-cell and spatial transcriptomics have generated vast atlases of cellular states, yet these data are almost exclusively used for analysis rather than generation. Here we introduce the LLM-based model PGL, Portraying Gene Language, a framework that reframes single-cell transcriptomes as a language generation modeling problem. PGL represents each cell as a long sequence of gene–expression tokens and uses a large language model to synthesize complete single-cell RNA-seq profiles from metadata alone, such as tissue and disease context. PGL-generated cells recapitulate dataset-specific transcriptomic structure, align with known cancer subtype biology, and mix coherently with real single-cell datasets. Notably, generated cells can be used as effective references for spatial transcriptomics, enabling accurate cell-type mapping without matched single-cell atlases. By shifting single-cell modeling from representation learning to cell generation, PGL enables virtual cohort construction, hypothesis generation, and reference-on-demand analysis, positioning generative language models as foundational tools for *in silico* single-cell biology.

## INTRODUCTION

Single-cell and spatial transcriptomics have rapidly expanded in scale and scope, with millions of cells profiled across thousands of datasets spanning diverse tissues, diseases, and experimental conditions ^1, 2^. As the field matures, the central question is no longer whether single cells can be measured, but whether the accumulated knowledge can be used to generate new, biologically plausible single cells *in silico*^3^.

Transformer-based models have already demonstrated strong utility in single-cell analysis^4^. Frameworks such as scGPT, Geneformer, TranscriptFormer, and related architectures leverage attention mechanisms to learn cell embeddings that support downstream tasks including cell typing, perturbation response prediction, and representation learning^5–7^. In parallel, language model–driven approaches such as CELLama and C2S have explored sequence-based formulations of single-cell and spatial transcriptomics, reframing gene expression profiles as languages to embed cellular features^8, 9^. These studies provide important evidence that large language models (LLMs) can capture the complexity of cellular gene expression programs. Despite these advances, existing transformer and language-based models primarily treat single-cell data as an embedding or representation learning problem. Their outputs are typically low-dimensional vectors optimized for prediction or classification tasks^10^. As a result, the notion of cell generation—producing a new single cell *in silico* with thousands of genes expressed in a coherent and condition-consistent manner—has remained largely unexplored. Here, we propose a different framing, instead of representing a cell as a high-dimensional numeric vector, we ask: *what if a single cell could be represented as a long piece of text?* In this view, a cell is described as a sequence of gene symbols paired with discretized expression scales, transforming gene expression into a paragraph-length description. Generating a cell thus becomes a language generation problem.

This formulation allows us to directly exploit the strengths of modern LLMs: modeling long-range dependencies, generating coherent and structured sequences, and conditioning outputs on rich, structured inputs. Building on this idea, we introduce PGL, Portraying Gene Language, a generative framework that uses LLMs to synthesize single-cell RNA-seq profiles from cell-level metadata alone. Given information such as tissue, disease context, or experimental condition, PGL autoregressively generates a long gene–expression sequence between special tokens, which is subsequently parsed back into a numeric gene expression matrix. We demonstrate that virtually generated single cell data recapitulate key statistical and biological properties of real data. Our results position PGL as a step toward truly generative single-cell biology, where virtual cells can be synthesized, compared, and analyzed using standard single-cell and spatial omics workflows.

## RESULTS

### Language-Based Representation and Generation of Single-Cell Transcriptomes

The model was based on a LLM capable of generating long-form text conditioned on designated metadata, including tissue of origin, disease status, and perturbation status. Given this metadata as input, the LLM generated textual representations of cellular states, and the model was trained to reconstruct sentences derived from ground-truth texts. The ground-truth texts were generated from single-cell RNA sequencing (scRNA-seq) datasets, in which each cell’s gene expression profile was represented as a sentence. Specifically, gene expression levels were discretized into quantized categories—Very High, High, Mid, Low, and Absent—and each gene was mapped to its corresponding expression grade. For each cell, sentences were constructed using a combination of top-expressed genes, marker gene sets identified from the scRNA-seq data, and randomly sampled genes. These sentence representations served as ground-truth texts for training the text-generation model (**Fig. 1a**).

**Figure 1.**
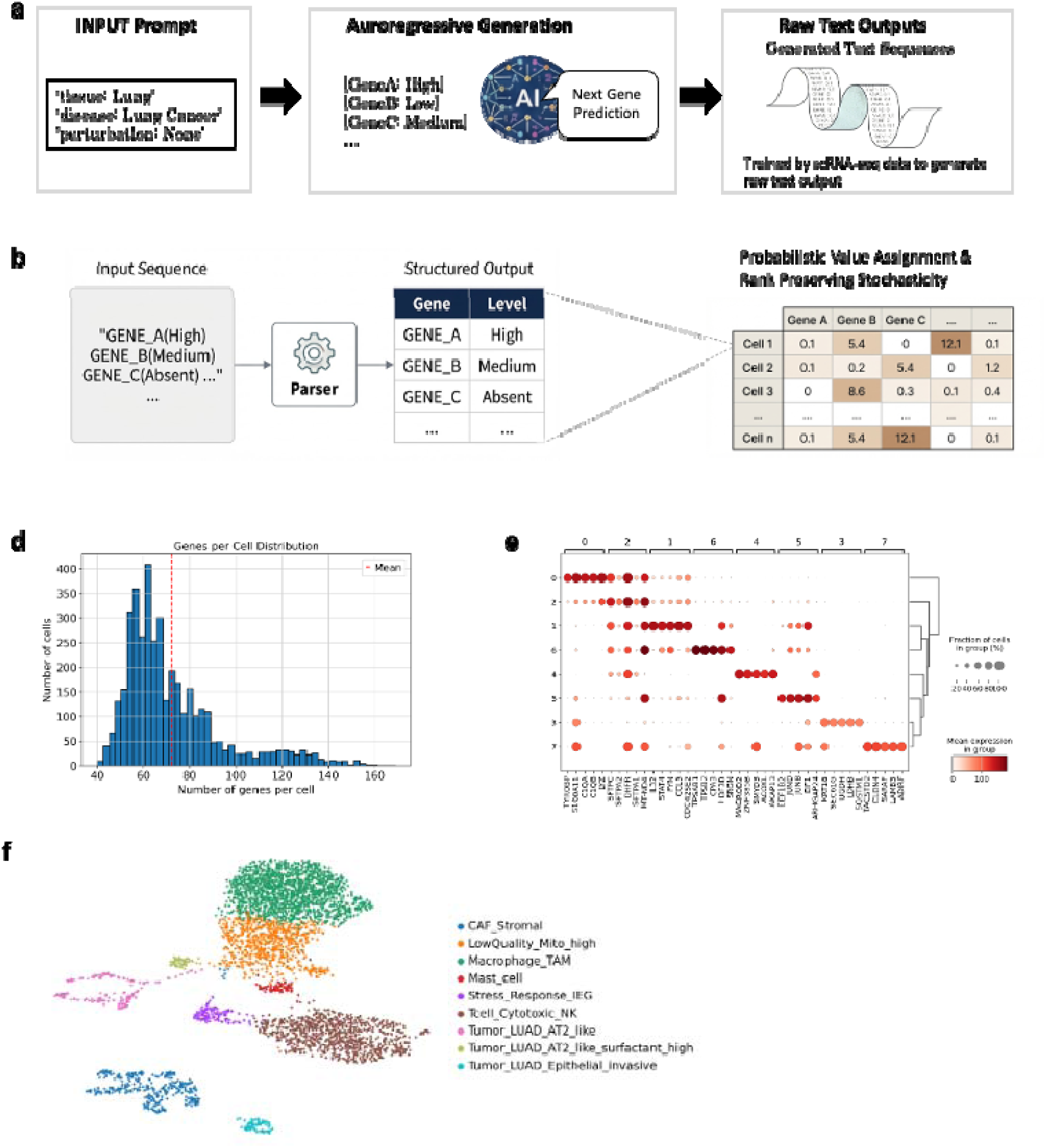
Language-based representation and generation of single-cell transcriptomes. a, Schematic overview of the PGL framework. Single-cell RNA-seq profiles are converted into sentence-like representations, where each cell is represented as an ordered sequence of gene–expression tokens with discretized expression levels (Very High, High, Mid, Low, Absent). Sentences are constructed using a combination of dataset-level marker genes, cell-level highly expressed genes, and sampled genes, and are used as ground-truth texts for training a large language model conditioned on cell-level metadata. b, Autoregressive generation and parsing pipeline. Given metadata alone, the trained PGL model generates gene–expression sentences, which are subsequently parsed back into numeric cell-by-gene expression matrices, enabling downstream single-cell analysis using standard workflows. c, Distribution of the number of genes per cell after parsing PGL-generated sequences into expression matrices. The generated lung adenocarcinoma cells show a median of 72.3 genes per cell. d, Expression of canonical marker genes illustrating biologically meaningful transcriptional patterns consistent with known cell identities. e, UMAP embedding of PGL-generated lung adenocarcinoma cells showing coherent clustering structure, with clusters annotated to cell types based on marker gene expression.

After training the PGL model on a general-purpose text-based language model (Qwen-0.6B), the model generated sentences in an autoregressive manner^11^. These generated texts were subsequently parsed into cell-by-gene expression matrices (**Fig. 1b**), enabling their interpretation as scRNA-seq data generated by the LLM^12^.

We evaluated whether LLM-generated scRNA-seq data exhibit global properties comparable to those of real scRNA-seq datasets. Using the PGL model after training from 14M tumor cells from 2119 scRNA-seq datasets, we generated 4,000 cells with a maximum token length of 8,192 from lung adenocarcinoma, corresponding to an average of approximately 72.3 genes per cell (**Fig. 1c**). Leiden clustering partitioned the generated cells into distinct clusters, and canonical marker genes enabled robust annotation of these clusters to known cell types (**Fig. 1d,e**), indicating coherent cellular identities within the generated data.

### Recapitulation of Dataset-Specific Transcriptomic Structure by LLMs

We assessed whether scRNA-seq data generated from metadata information recapitulate dataset-specific transcriptomic characteristics. We measured similarity between the generated data and real scRNA-seq datasets sampled from 10 datasets with matching metadata and 10 datasets with mismatched metadata from different tissue origins and/or different diseases^2^.

We first evaluated global transcriptomic similarity by computing Spearman rank correlations between gene expression distributions of generated and real datasets. Across same metadata comparisons (lung adenocarcinoma data, synthetic vs real), generated data exhibited consistently higher correlations compared to different metadata datasets. The mean Spearman correlation was significantly higher for same metadata datasets than for different metadata controls (0.47±0.04 vs. 0.34±0.05; p < 0.001), indicating that generated scRNA-seq data more closely resemble real scRNA-seq data derived from the same biological context (**Fig. 2a, Supplementary Fig. 1a** showed examples of correlation across genes).

**Figure 2.**
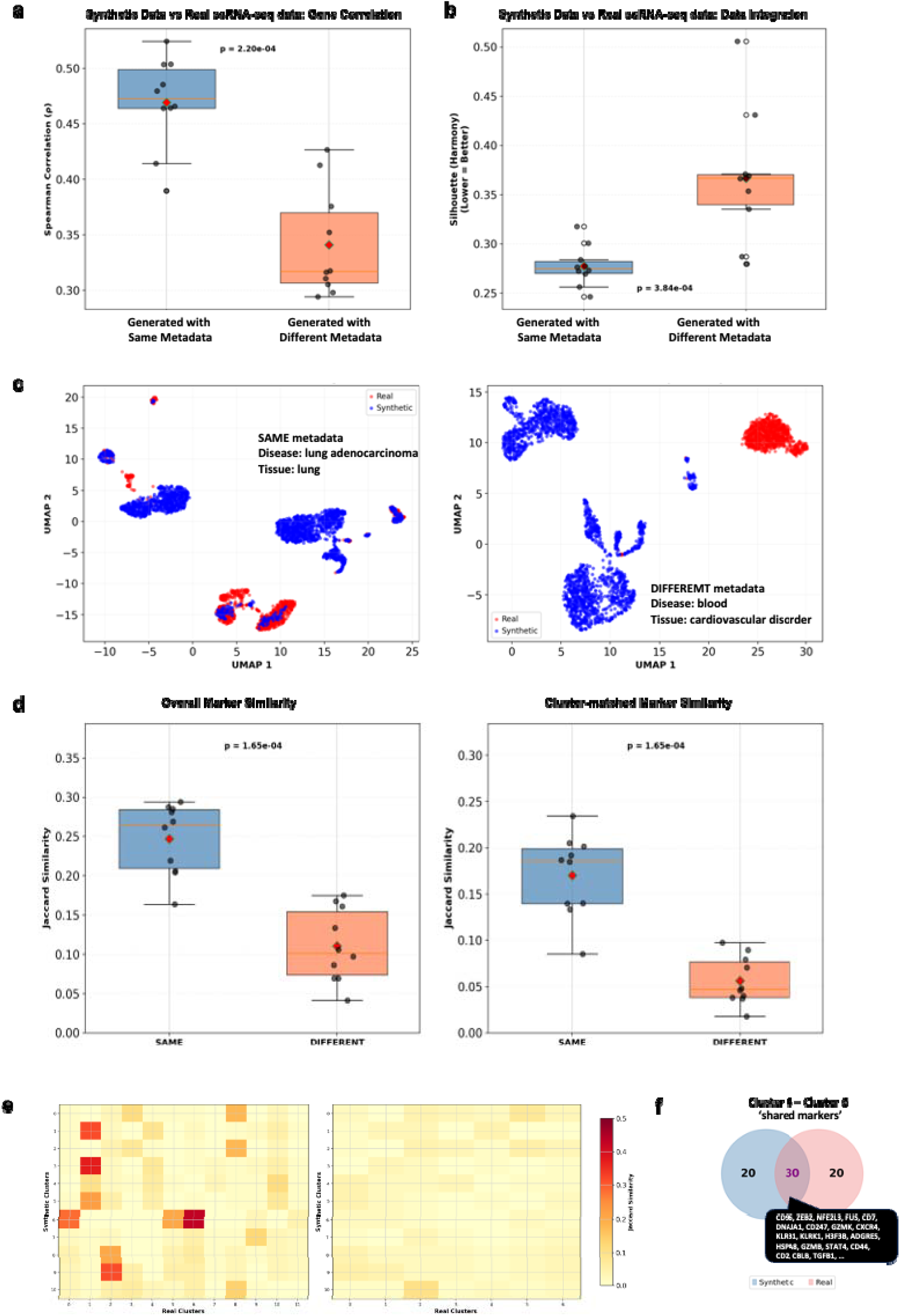
Recapitulation of dataset-specific transcriptomic structure by LLM-generated cells. a, Global transcriptomic similarity between generated scRNA-seq data and real datasets quantified by Spearman rank correlation of gene-wise expression distributions. Generated lung adenocarcinoma cells show significantly higher similarity to real datasets with matching metadata than to mismatched tissue or disease datasets. b, Single-cell–level similarity assessed by silhouette scores following integration of generated and real datasets, with batch correction. Lower silhouette scores indicate improved mixing between generated and real cells for matched metadata compared to mismatched controls. c, Representative UMAP embeddings of integrated generated and real scRNA-seq datasets, illustrating improved mixing under matched metadata conditions. d, Marker gene overlap quantified by overall and cluster-matched Jaccard similarity. Both metrics show significantly higher overlap between generated and real datasets sharing the same metadata. e, Heatmaps of pairwise Jaccard indices between clusters of generated and real datasets, demonstrating higher marker gene overlap across multiple clusters under matched metadata conditions. f, Representative example of shared cluster-specific marker genes between generated and real lung adenocarcinoma scRNA-seq datasets, illustrating concordant transcriptional programs.

To assess transcriptomic similarity at the single-cell level, we integrated LLM-generated and real scRNA-seq datasets and quantified their degree of mixing using silhouette scores^13^. Both with and without batch correction^14^, integrations with real datasets sharing the same metadata yielded significantly lower silhouette scores compared to integrations with datasets of mismatched metadata, indicating improved mixing and reduced separability between generated and real cells. After batch correction, integrations with same metadata datasets resulted in a mean silhouette score of 0.28 ± 0.02 (n = 10), whereas integrations with different metadata datasets exhibited substantially higher scores (0.37 ± 0.06, n = 10; Mann–Whitney U test, P = 3.8 × 10LL) (**Fig. 2b**). A similar trend was observed without batch correction, with same metadata integrations showing lower silhouette scores (0.31 ± 0.02, n = 10) compared to different metadata integrations (0.37 ± 0.05, n = 10; Mann–Whitney U test, P = 2.9 × 10L³) (**Supplementary Fig. 1b**). These results demonstrate that generated cells are more transcriptionally similar to real cells derived from the same tissue and disease context than to unrelated datasets. The examples of integrated UMAP results are represented in **Fig. 2c**.

We further quantified overlap in transcriptomic marker features between generated and real scRNA-seq data using Jaccard similarity metrics. Following clustering of both synthetic and real datasets, cluster-specific marker genes were identified and compared using two complementary approaches: *overall marker similarity* and *cluster-matched marker similarity*. Overall similarity was computed by comparing the union of marker genes across datasets, whereas cluster-matched similarity was defined as the mean of the maximum Jaccard index obtained for each cluster against all clusters in the counterpart dataset. The overall Jaccard index was significantly higher for same metadata comparisons than for different metadata controls (0.25 ± 0.04 vs. 0.11 ± 0.04; Mann–Whitney U test, P = 1.7 × 10LL), indicating greater global overlap in marker gene composition when generated data were compared to real datasets sharing the same metadata. Similarly, cluster-matched Jaccard similarity was markedly higher in same metadata integrations (0.17 ± 0.04) compared to different metadata integrations (0.056 ± 0.025; Mann–Whitney U test, P = 1.7 × 10LL), reflecting improved correspondence between transcriptionally matched clusters (**Fig. 2d**). Representative examples of cluster-level marker gene similarities are visualized in **Fig. 2e**. Heatmaps of pairwise Jaccard indices across clusters demonstrate that datasets sharing the same metadata exhibit consistently higher marker overlap across multiple clusters, whereas datasets with different metadata show substantially reduced overlap. An illustrative example of shared marker genes corresponding to the cluster pairs highlighted in **Fig. 2e** is shown in **Fig. 2f**, where synthetic and real lung adenocarcinoma scRNA-seq datasets display substantial concordance in cluster-specific marker genes.

### Comparative analysis of generated data recapitulates known cancer biology

We next examined whether comparisons between independently generated scRNA-seq datasets could recover established biological differences and enable tumor subtype–specific marker discovery. As a case study, we contrasted hormone receptor–positive (HR-positive) and triple-negative breast cancer (TNBC), two subtypes with well-characterized lineage and transcriptional differences. Using the trained PGL model, we generated 10,000 synthetic cells for each subtype. The two datasets were integrated and visualized using UMAP, with expression of canonical luminal markers (ESR1, KRT8, KRT18), basal markers (KRT5, KRT14, EGFR), and proliferation markers (MKI67, TOP2A, CDK1) overlaid on the embedding (**Fig. 3a**). Marker expression patterns recapitulated known subtype-specific biology, with luminal markers enriched in HR-positive cells and basal markers predominating in TNBC^15, 16^.

**Figure 3.**
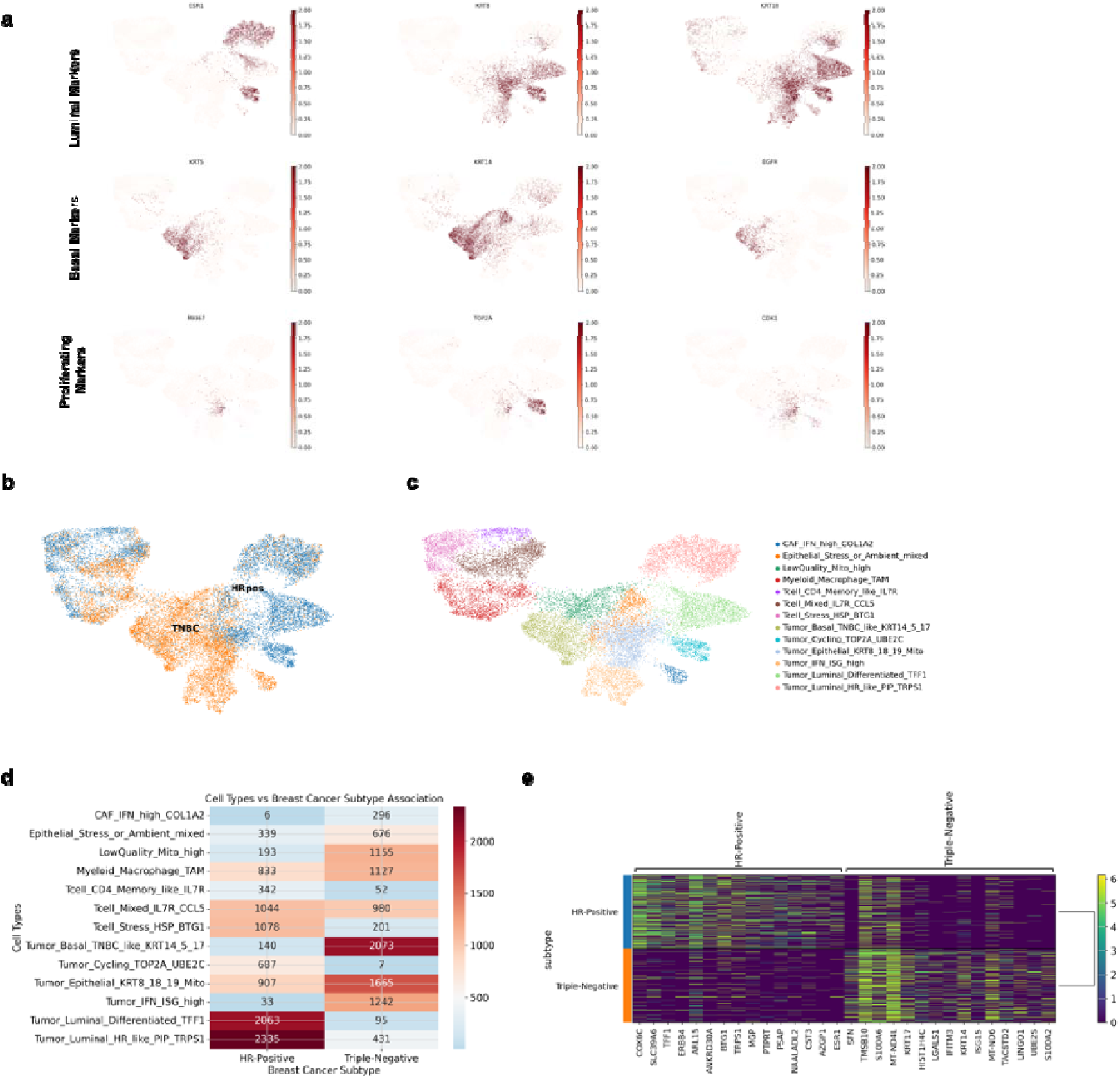
Comparative analysis of independently generated breast cancer subtypes. a, UMAP embedding of independently generated hormone receptor–positive (HR-positive) and triple-negative breast cancer (TNBC) cells (10,000 cells per subtype), with expression of canonical luminal (ESR1, KRT8, KRT18), basal (KRT5, KRT14, EGFR), and proliferation (MKI67, TOP2A, CDK1) markers overlaid. b, UMAP colored by breast cancer subtype, showing clear separation between HR-positive and TNBC synthetic cohorts. c, UMAP colored by marker-defined cell types, illustrating coherent subtype-associated transcriptional structure. d, Comparison of cell type proportions between HR-positive and TNBC generated datasets, revealing subtype-specific compositional differences. e, Differential expression analysis between synthetic HR-positive and TNBC cells, highlighting top subtype-specific marker genes.

UMAP embeddings colored by breast cancer subtype and marker-defined cell types further demonstrated clear separation and coherent subtype-associated transcriptional structure (**Fig. 3b,c; Supplementary Fig. 2a**). Consistent with these observations, curated gene sets exhibited subtype-specific expression shifts, with luminal gene sets significantly enriched in HR-positive breast cancer and basal gene sets enriched in TNBC (**Supplementary Fig. 2b**), confirming that the generated data align with established knowledge-based marker signatures. We additionally compared cell type proportions between the two generated subtypes, revealing distinct compositional differences consistent with known biological characteristics of HR-positive and TNBC tumors (**Fig. 3d**). Finally, differential expression analysis between the synthetic HR-positive and TNBC datasets identified subtype-specific marker genes, with top differentially expressed genes summarized in **Fig. 3e**.

### Model-generated cells enable spatial mapping in spatial transcriptomics data

We next evaluated whether generated single-cell transcriptomic profiles could serve as effective references for spatial transcriptomics analysis. Using Visium spatial gene expression datasets, we mapped generated cells onto spatial coordinates via standard cell type transfer and deconvolution pipelines. The generated cells exhibited coherent spatial localization patterns that aligned with known tissue architecture, including tumor–stroma boundaries.

As a primary case study, we used PGL model–generated prostate cancer scRNA-seq data alongside a real prostate cancer atlas dataset^17^. After clustering each dataset independently, broad cell types—epithelial, fibroblast, endothelial, myeloid, and lymphocyte populations—were annotated based on canonical marker genes (**Fig. 4a,b**; Heatmaps for markers in **Supplementary Fig. 3**). These annotated cell types were then transferred to prostate cancer Visium data using an optimal transport–based deconvolution method (TACCO)^18^. Spatial cell type maps derived from synthetic and real reference datasets showed highly concordant spatial distributions (**Fig. 4c**). To quantitatively assess agreement between the two approaches, we performed spot-wise correlation analysis of inferred cell type proportions. This analysis revealed significant positive correlations across all cell types between synthetic-based and real-based mappings (**Fig. 4d**), indicating that generated scRNA-seq data provide spatial annotations comparable to those obtained using real reference datasets.

**Figure 4.**
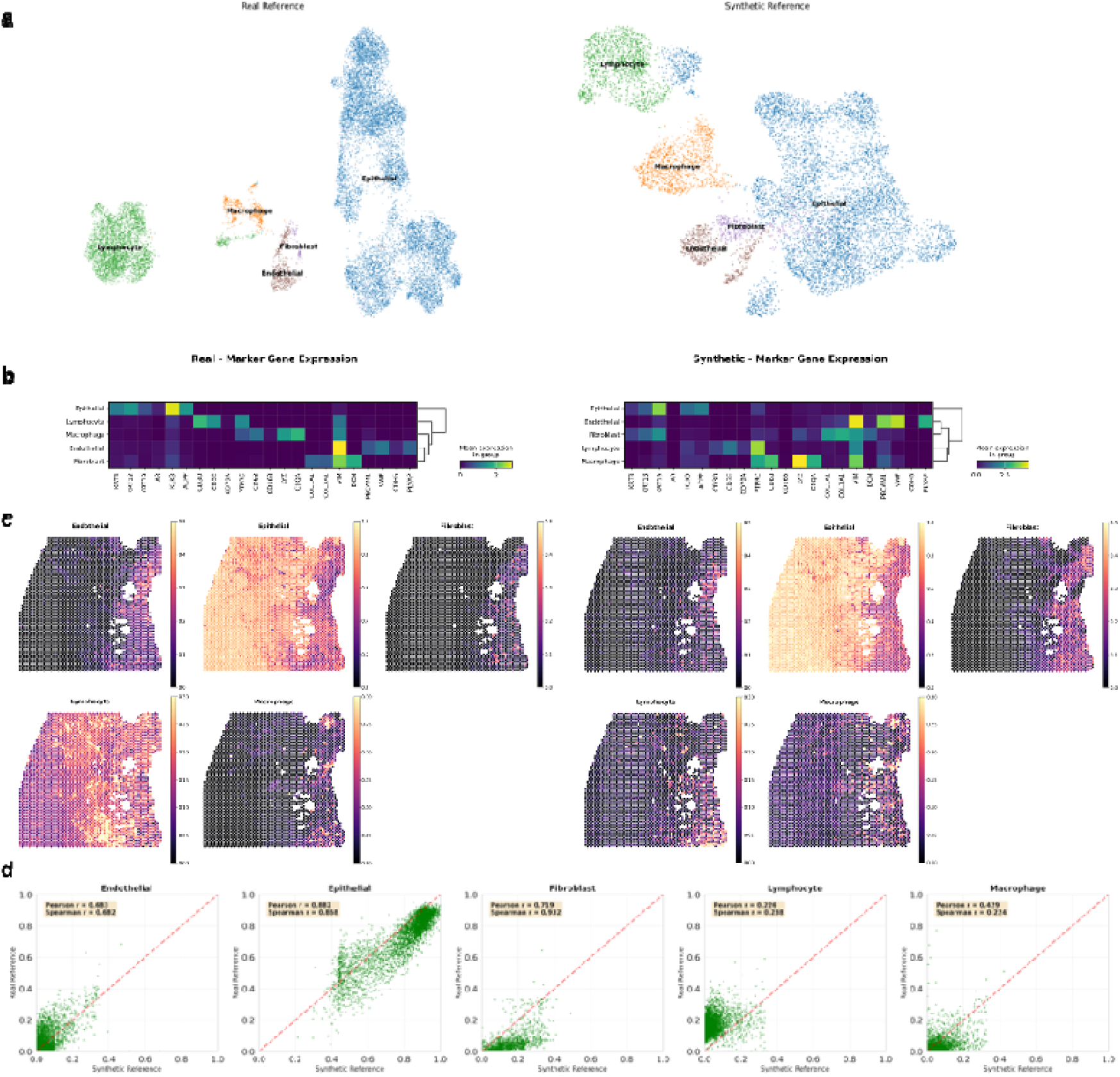
Model-generated cells as references for spatial transcriptomics. a, UMAP embeddings of real (left) and PGL-generated (right) prostate cancer scRNA-seq datasets. b, Clustering and broad cell type annotation of PGL-generated prostate cancer scRNA-seq data based on canonical marker genes. c, Spatial cell type maps inferred for prostate cancer Visium data using synthetic and real scRNA-seq references with optimal transport–based deconvolution, showing highly concordant spatial distributions. d, Spot-wise correlation of inferred cell type proportions between synthetic-based and real-based mappings across cell types, demonstrating strong agreement.

To verify that these results were not specific to a single transfer method, we repeated the analysis using an independent deep learning–based deconvolution approach (CellDART)^19^. This alternative method produced highly similar spatial cell type distributions and similarly strong positive correlations between synthetic- and real-based mappings across all cell types (**Supplementary Fig. 4a,b**). Finally, we extended this evaluation to a higher-resolution spatial transcriptomics platform using Visium HD breast cancer data with 16-μm spatial bins. Consistent with the prostate cancer analyses, synthetic-based and real-based cell type mappings exhibited significant positive correlations across inferred cell types (**Supplementary Fig. 5**), demonstrating robustness across tissue types and spatial resolutions. Together, these results demonstrate that PGL-generated single-cell transcriptomic profiles can serve as reliable reference datasets for spatial transcriptomics, enabling accurate cell type mapping across diverse tissues, platforms, and analytical frameworks.

## DISCUSSION

The advent of LLM has transformed our ability to model complex, long-range structured sequences, extending from natural language to biological information systems^20^. In this study, we introduce PGL, a generative framework that reframes single-cell transcriptomics not as fixed numerical vectors, but as a structured linguistic process. By representing a cell as a paragraph-length sequence of gene–expression tokens, PGL leverages the intrinsic strengths of decoder-only, autoregressive language models, their capacity to model long-context dependencies, maintain global coherence, and generate structured outputs token by token^21^. This formulation shifts single-cell modeling from representation learning toward explicit *in silico* cell generation, enabling the synthesis of complete transcriptomes conditioned on biological context. Our results demonstrate that PGL-generated cells are biologically coherent entities that recapitulate dataset-specific transcriptional programs across diverse tissues and disease states.

Existing transformer-based models in the single-cell field, such as scGPT and Geneformer, have primarily focused on cell embedding for classification and perturbation prediction^5–7^. While highly effective, these approaches typically represent each cell as a point in a latent space optimized for discrimination rather than generation. PGL departs from this embedding-centric paradigm by demonstrating that a decoder-only language model can autoregressively synthesize full, high-dimensional transcriptomes from minimal cell-level metadata. This generative capability implies that the statistical regularities of cellular gene expression—the combinatorial rules governing gene co-expression across biological contexts—are implicitly encoded within the model parameters. The ability of refined LLM to generate cells that mix with real datasets in UMAP space, even maintaining dataset-specific marker signatures (as evidenced by Jaccard similarities), underscores its potential as a “digital twin” generator for single-cell biology. This observation that PGL-generated cells integrate with real datasets in low-dimensional embeddings and preserve dataset-specific marker signatures further supports the view that generative LLMs can function as digital counterparts of biological cells, rather than merely as feature extractors for downstream analyses^22^.

The utility of PGL extends beyond data replication toward *in silico* hypothesis generation. In comparative analyses of independently generated breast cancer subtypes, PGL recapitulated well-established transcriptional differences between hormone receptor–positive and triple-negative breast cancer, recovering canonical luminal and basal marker programs^15,16^. These results indicate that the model preserves subtype-defining gene expression structure when generating cells conditioned only on disease metadata. Beyond expected lineage markers, PGL also highlighted subtype-associated genes that are consistent with prior clinical and translational observations. For example, regarding LGALS1(Galectin-1), recent studies have identified LGALS1 as an independent prognosticator of poor survival in TNBC^23^. It is known to promote an immunosuppressive “shield” in the tumor stroma, facilitating immune escape and driving the epithelial-mesenchymal transition^24^. In addition, IFITM3, while traditionally known for its antiviral roles, IFITM3 has emerged as an oncogenic driver in invasive breast cancers ^25^. Its high expression in PGL-generated TNBC cohorts reflects its established role in cell cycle progression and “immuno-hot” tumor microenvironments, where it correlates with high-grade malignancy and poor clinical outcomes^26^. Although these observations do not constitute biological validation or the discovery of new markers, they suggest that the generative model captures higher-order transcriptional programs that reflect known disease mechanisms. In this context, PGL may serve as a complementary exploratory tool, enabling systematic comparison of disease states and prioritization of candidate genes for downstream experimental investigation. The fact that PGL can autonomously “rediscover” such markers suggests its utility as a hypothesis-generation engine. For researchers working with rare diseases or limited clinical samples, PGL can generate pseudo-single-cell cohorts to identify potential therapeutic targets or biomarkers before moving to expensive in vitro or in vivo validation.

One of the most practical applications demonstrated in this study is the use of PGL-generated cells as reference datasets for ST analysis. Accurate cell-type mapping in ST typically depends on the availability of a high-quality, well-matched scRNA-seq atlas derived from the same tissue, disease state, and experimental context^27^. In practice, such matched references are often unavailable, incomplete, or prohibitively expensive to generate, particularly for rare tissues, specific disease subtypes, or clinical samples with limited material^28^. Even when scRNA-seq data exist, differences in platform, sample preparation, or cohort composition can introduce substantial variability, complicating their use as reference atlases for spatial deconvolution. PGL addresses these limitations by enabling the synthesis of reference cell populations that are explicitly conditioned on the metadata of the spatial sample. Rather than relying on a single experimental atlas, PGL can generate reference-on-demand scRNA-seq profiles tailored to the tissue and disease context under study. Using several deconvolution methods, we show that synthetic references yield spatial cell-type maps and spot-wise cell-type proportions that closely match those obtained using real scRNA-seq atlases. Importantly, this concordance is observed across multiple cell types, analytical frameworks, and spatial resolutions. At the same time, synthetic references are not a replacement for experimentally derived atlases. Generated cells do not capture patient-specific genetic variation, rare or unexpected cell states absent from the training data, or technical artifacts unique to a given experiment. Instead, their strength lies in providing a biologically informed baseline reference that encodes typical transcriptional programs for a given context. In this role, PGL-generated references can substantially lower the barrier to ST analysis, enabling exploratory cell-type mapping, comparative spatial studies, and hypothesis generation in settings where matched single-cell data are unavailable. As spatial transcriptomics continues to expand into clinical and translational applications, such flexible, metadata-driven reference construction may represent a pragmatic complement to experimental atlas generation.

As single-cell datasets continue to grow, PGL offers a scalable solution for data augmentation. By synthesizing diverse and condition-consistent cells, PGL can provide the “pre-training” data necessary to make downstream classifiers more robust, particularly for underrepresented populations or various cell types. However, the current implementation of PGL has several limitations. First, discretizing gene expression into a small number of ordinal categories facilitates the formulation of transcriptomes as a language modeling problem, but inevitably compresses the continuous dynamic range of raw count data. Although percentile-based binning preserves relative expression rankings within each cell, fine-grained quantitative differences between genes and subtle expression shifts across conditions may be partially lost. Future extensions could incorporate finer-grained discretization schemes or hybrid formulations that combine categorical tokens with continuous-valued outputs. Second, the generation capacity of PGL is constrained by the maximum context length of current LLMs. Because each gene–expression pair consumes multiple tokens, only a subset of genes can be generated per cell within a fixed token budget, resulting in sparsity in the reconstructed expression matrix. While hierarchical gene sampling mitigates this issue during training, incomplete gene coverage remains a limitation during inference. Advances in long-context LLM architectures, increased maximum token lengths, and scaling to larger backbone models are expected to substantially alleviate this constraint, enabling denser and more comprehensive transcriptome generation. Additionally, iterative or multi-pass generation strategies may further expand effective gene coverage without exceeding context limits. Together, these limitations highlight that PGL represents an early step toward generative single-cell modeling, with clear avenues for improvement as language model architectures and training paradigms continue to evolve. Future iterations could integrate more granular scales or multi-omic metadata to create a more holistic “cell language.” In conclusion, PGL positions generative LLMs as a foundational tool for the next era of biology. By moving from observing cells to synthesizing them, we move closer to a predictive framework where the response of a tissue to a novel disease or therapeutic can be modeled entirely in silico, accelerating the pace of biological discovery.

## METHODS

### Data curation and preprocessing

PGL model is a generative framework that models single-cell gene expression as a language generation problem. The core idea is to represent a single cell as a long sequence of gene–expression tokens and to train a LLM to autoregressively generate such sequences conditioned solely on cell-level metadata.

Single-cell RNA-seq data were curated from the scBaseCount repository^2^, focusing on cancer datasets profiled using droplet-based platforms. After quality control and firstly selecting the focus on the scRNA-seq data from tumors, the final training corpus comprised approximately 14 million cells derived from 2,118 independent datasets, including lung, breast, colorectal, pancreatic, and others (**Supplementary Table 1**).

Each dataset included raw count matrices and associated metadata. Cell-level metadata were standardized across datasets to include tissue, organ, disease type, and experimental or perturbation condition when available.

### Quality control

Cells with fewer than 200 detected genes were removed. No explicit upper bound was placed on gene counts in order to preserve highly complex malignant or activated immune cells. Genes expressed in fewer than 0.1% of cells within a dataset were excluded to reduce noise from ultra-rare transcripts. Raw counts were retained for downstream discretization. No global normalization or log-transformation was applied prior to gene binning, ensuring that relative expression rankings were computed within individual cells.

### Gene panel harmonization

To ensure compatibility with spatial transcriptomics platforms, gene sets were filtered to predefined panels depending on the training configuration. The model training used Visium-compatible gene panels, predefined by 10X genomics (https://www.10xgenomics.com/support/spatial-gene-and-protein-expression/documentation/steps/probe-sets), to reduce the token size during the training and inference process, considering these gene panels cover various gene sets to characterize disease and tissue status. Gene identifiers were mapped to standardized gene symbols prior to filtering.

### Marker-aware gene selection

For each dataset, unsupervised clustering was performed to identify dataset-specific marker genes. Cells were subsampled if necessary, normalized, and log-transformed for clustering only. Highly variable genes were selected, followed by dimensionality reduction and Leiden community detection. For each cluster, differentially expressed genes were identified using a rank-based statistical test, and top-ranking genes were aggregated to define a dataset-level marker gene list.

### Hierarchical gene sampling per cell

For each training cell, genes were selected using a three-tier hierarchy:

1. Dataset-level marker genes, capturing dominant cell types and transcriptional programs.
2. Cell-level highly expressed genes, capturing transient or state-specific signals unique to the cell.
3. Randomly sampled genes from the remaining panel, ensuring broad gene coverage and preventing overfitting.

This strategy resulted in approximately 200 genes per training example, enabling efficient training while encouraging generalization beyond the observed gene subsets.

### Expression discretization and tokenization

Continuous gene expression values were discretized into five ordinal categories: Absent, Low, Medium, High, and Very High. Binning was performed on a per-cell basis using percentile thresholds computed from expressed genes only. This approach normalizes for library size variation and emphasizes relative expression patterns within each cell.

Selected genes were ordered by decreasing expression level and serialized into a text sequence of the form “GENE(ExpressionLevel)”. Each sequence was wrapped between special tokens <START_GEN> and <END_GEN>. A single generated cell typically corresponds to 8,000–16,000 tokens, depending on the number of genes generated during inference.

### Overview of conditional single-cell generation

We model scRNA-seq generation as conditional sequence modeling. Given cell-level metadata *c* (tissue, organ, disease status, perturbation), the model generates an ordered sequence of gene–expression tokens of the form

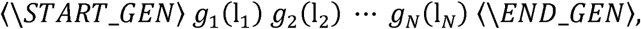

where *g_i_* is a gene symbol and l_*i*_ ϵ (Absent, Low, Med, High, Very High) is a discretized expression level. The conditional likelihood factorization follows standard causal language modeling:

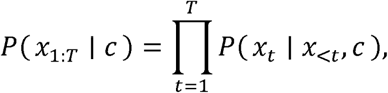

where *x*_1:*T*_ are tokens representing the metadata prompt and the gene–level sequence.

### Model architecture and adaptation

A pretrained decoder-only transformer language model was used as the backbone. The model supports long-context autoregressive generation and was selected to balance representational capacity with computational efficiency. In this experiment, we used Qwen3-0.6B as the base decoder-only transformer. New embeddings were initialized by averaging embeddings of semantically related tokens from the base vocabulary.

Low-Rank Adaptation (LoRA) was applied to all attention and feed-forward projection layers^29^. Only LoRA parameters were updated during training, while the base model weights remained frozen. This approach enables efficient scaling to millions of training examples while mitigating overfitting and memory overhead. Cell metadata were provided as structured natural-language prompts preceding the gene-expression sequence, allowing the model to condition generation on biological context. LoRA applied to attention and MLP projection layers. For a frozen base weight matrix *W*_0_ ϵ *R^d×k^*, LoRA introduces a low-rank update:

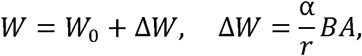

where *B* ϵ *R^d×r^* and *A* ϵ *R^r×k^* are trainable matrices, r is the rank, and α is a scaling hyperparameter. We used r=32, a =64, and LoRA dropout 0.05.

### Training procedure

The model was trained using a causal language modeling objective to maximize the likelihood of the gene-expression token sequence conditioned on metadata and preceding tokens. Loss was computed only on gene-expression tokens, excluding prompt tokens. Training prompts were dynamically regenerated at each epoch with different random gene selections to improve robustness and prevent memorization of specific gene combinations. We trained the model using causal language modeling with teacher forcing. F^12^ or token sequence *x*_1:*T*_ (including the gene expression segment), the loss is:

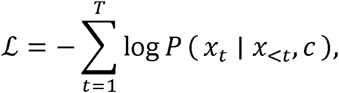

computed only on target tokens corresponding to the gene–level sequence (prompt tokens were masked from loss).

Multiple training scales were explored, including pilot experiments on a small number of datasets and large-scale training across thousands of samples. Hyperparameters such as learning rates and batch sizes were selected empirically and are adjustable within the same framework.

### Autoregressive generation and post-processing

At inference, the model receives only cell-level metadata as input and autoregressively generates a gene-expression sequence between <START_GEN> and <END_GEN> tokens. Sampling-based decoding was used to promote diversity while preserving biological plausibility.

Generated sequences were parsed to extract gene–expression pairs. Discrete expression levels were stochastically mapped back to continuous numeric values, yielding a full gene-by-cell expression matrix. Multiple generated cells were aggregated into standard AnnData objects for downstream analysis^12^.

### Selection of matched and mismatched real scRNA-seq datasets

To evaluate whether metadata-conditioned generation recapitulates dataset-specific transcriptomic structure, we assembled two comparison groups of real scRNA-seq datasets: (i) matched metadata datasets (same tissue and disease context as the generated data; n=10) and (ii) mismatched metadata datasets originating from different tissue and/or disease contexts (n=10) from scBaseCount (**Supplementary Table 2**). Unless otherwise specified, comparisons were performed between the generated dataset and each real dataset independently, yielding a set of similarity statistics per group.

### Global similarity via gene-wise distribution correlation

We quantified global similarity between a generated dataset and a real dataset by comparing gene expression distributions aggregated at the gene level. For each gene *g*, we computed the sum of expression distribution across cells (per-gene sum expression across cells; identical processing was applied to generated and real datasets). We then computed Spearman rank correlation between the resulting gene-wise vectors. Group-level comparisons (matched vs mismatched) were performed on the per-dataset *rho* values, and Mann Whitney test was conducted for the comparison.

### Single-cell level integration and mixing quantified by silhouette score

To assess similarity at the single-cell level, we integrated generated and real scRNA-seq datasets into a shared embedding and quantified mixing using silhouette scores. Prior to integration, both datasets were processed using a consistent single-cell analysis pipeline, including normalization, highly variable gene selection, and PCA. Integration was performed in two modes: with batch correction and without batch correction. For batch-corrected integration, we applied Harmony on the joint PCA space using dataset identity (generated vs real) as the batch covariate^14^. For uncorrected integration, UMAP was computed directly from the joint PCA space. To quantify mixing, we computed silhouette scores with respect to dataset labels (generated vs real). For each cell *i*, the silhouette coefficient is:

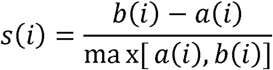

where *a(i)* is the mean distance from cell *i* to other cells in the same dataset label and *b(i)* is the minimum mean distance to cells in the other dataset label. We summarized dataset-level mixing by averaging *s(i)* across cells for each integrated pair. Lower silhouette values indicate improved mixing (reduced separability) between generated and real cells.

### Marker gene overlap quantified by Jaccard similarity

To compare transcriptomic features at the level of marker genes, we clustered generated and real datasets independently and computed cluster-specific marker genes using differential expression analysis (one-vs-rest). For each dataset, marker gene sets were derived per cluster; downstream comparisons were performed using two complementary Jaccard metrics^30^.

Overall marker similarity: Let *M*_syn_ and *M*_real_ denote the union of marker genes across all clusters in the synthetic and real datasets, respectively. The overall Jaccard similarity is:

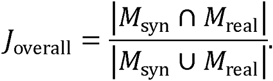

Cluster-matched similarity: Let *M*^(*k*)^_syn_ be the marker set for synthetic cluster k and *M*^(*j*)^_real_ for real cluster j. For each synthetic cluster k, we computed the maximum Jaccard overlap across all real clusters:

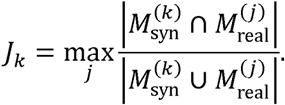

The cluster-matched similarity was then computed as the mean across synthetic clusters:

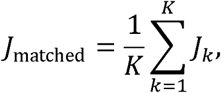

where K is the number of synthetic clusters.

### Integration and visualization of subtype structure

The synthetic datasets (HR-positive and TNBC) were integrated using a standard Scanpy workflow. Briefly, counts were normalized, log-transformed, and highly variable genes were selected prior to PCA and neighborhood graph construction. UMAP was computed from the integrated neighbor graph. To visualize biological signal, we overlaid canonical marker gene expression on the UMAP embedding, including luminal markers (e.g., ESR1, KRT8, KRT18), basal markers (KRT5, KRT14, EGFR), and proliferation markers (MKI67, TOP2A, CDK1).

### Marker-based cell type assignment

Cell populations were annotated using marker-based rules derived from canonical lineage markers. Cells were assigned to broad cell states/types by thresholding relative marker expression within the embedding (e.g., luminal-like, basal-like, proliferating), and subtype-level distributions were visualized by coloring UMAP by subtype and marker-defined cell type.

### Differential expression analysis between subtypes

We performed differential expression testing between clusters of subtypes of generated scRNA-seq data. Genes were ranked by effect size and statistical significance (Wilcoxon rank-sum test), and top markers were summarized for visualization.

### Reference datasets for spatial mapping

To evaluate whether generated scRNA-seq profiles can serve as references for spatial transcriptomics, we used: (i) a synthetic prostate cancer scRNA-seq reference generated by the PGL model, and (ii) a real prostate cancer atlas scRNA-seq reference^17^. Both references were processed independently with clustering and marker-based broad cell type annotation. For breast cancer, we also used a real breast cancer atlas reference^16^. Cell types included epithelial, fibroblast, endothelial, myeloid, and lymphocyte populations. For each reference, clustering was performed using a standard single-cell workflow. Broad cell types were assigned based on canonical marker genes. Cluster-to-cell-type mapping was performed by inspecting marker enrichment per cluster; annotated cell types were then used as labels for downstream transfer.

### Spatial transcriptomics datasets

We analyzed prostate cancer Visium spatial gene expression data (https://www.10xgenomics.com/datasets/human-prostate-cancer-adenocarcinoma-with-invasive-carcinoma-ffpe-1-standard-1-3-0) and breast cancer VisiumHD spatial gene expression data (https://www.10xgenomics.com/datasets/visium-hd-cytassist-gene-expression-human-breast-cancer-fixed-frozen). Spots were filtered using standard QC, and gene identifiers were harmonized between spatial and reference datasets by intersecting gene symbols.

### Cell type transfer to ST data

We performed spatial cell type mapping using TACCO, an optimal transport–based deconvolution/label transfer framework^18^. We ran TACCO with default parameters separately using the synthetic reference and the real reference, producing two spot-by-cell-type proportion maps for direct comparison. To assess method robustness, we repeated spatial mapping using CellDART^19^, a deep learning–based deconvolution approach. CellDART was run with identical input preprocessing and cell type labels, and synthetic-vs real-reference mapping outputs were compared.

### Agreement between synthetic- and real-reference mapping

To quantify concordance between synthetic- and real-reference spatial maps, we computed spot-wise correlation of inferred cell type proportions for each cell type. We computed correlation across spots using Pearson or Spearman correlation consistently across analyses. Statistical significance was assessed by testing the null hypothesis of zero correlation across spots for each cell type (two-sided test).

### Statistical testing

Group comparisons between matched and mismatched metadata were performed on per-dataset similarity metrics. For silhouette and Jaccard comparisons, we used a two-sided Mann–Whitney U test to compare matched vs mismatched distributions. For Spearman correlations, group differences were tested on the set of per-dataset *rho* values (two-sided test).

### Data Availability

All single-cell RNA-seq datasets used for training and evaluation were curated from the scBaseCount repository. Publicly available reference scRNA-seq atlases used for downstream validation include a prostate cancer single-cell atlas from Tuong et al. (Cell Reports 2021, 37, 110132; https://doi.org/10.1016/j.celrep.2021.110132) and a breast cancer single-cell atlas (Nat. Genet. 2021; https://www.nature.com/articles/s41588-021-00911-1). Spatial transcriptomics datasets were obtained from 10x Genomics public resources. Prostate cancer Visium data were downloaded from: https://www.10xgenomics.com/datasets/human-prostate-cancer-adenocarcinoma-with-invasive-carcinoma-ffpe-1-standard-1-3-0. Breast cancer Visium HD data (16-µm binning) were downloaded from: https://www.10xgenomics.com/datasets/visium-hd-cytassist-gene-expression-human-breast-cancer-fixed-frozen. All analyses used gene symbols harmonized by intersection between reference and spatial datasets after standard quality control filtering.

## Acknowledgment

This research was supported by the National Research Foundation of Korea (NRF-2020M3A9B6038086 and 2023R1A2C2006636).

## References

1. Program, C.C.S. et al. CZ CELLxGENE Discover: a single-cell data platform for scalable exploration, analysis and modeling of aggregated data. Nucleic acids research 53, D886–D900 (2025).

2. Youngblut, N.D. et al. scBaseCount: an AI agent-curated, uniformly processed, and continually expanding single cell data repository. bioRxiv, 2025.2002. 2027.640494 (2025).

3. Bunne, C. et al. How to build the virtual cell with artificial intelligence: Priorities and opportunities. Cell 187, 7045–7063 (2024).

4. Szałata, A. et al. Transformers in single-cell omics: a review and new perspectives. Nature methods 21, 1430–1443 (2024).

5. Cui, H. et al. scGPT: toward building a foundation model for single-cell multi-omics using generative AI. Nature Methods, 1–11 (2024).

6. Theodoris, C.V. et al. Transfer learning enables predictions in network biology. Nature 618, 616–624 (2023).

7. Pearce, J.D. et al. A Cross-Species Generative Cell Atlas Across 1.5 Billion Years of Evolution: The TranscriptFormer Single-cell Model. bioRxiv, 2025.2004. 2025.650731 (2025).

8. Park, J. et al. CELLama: foundation model for single cell and spatial transcriptomics by cell embedding leveraging language model abilities. Advanced Science, e13210 (2024).

9. Levine, D. et al. Cell2Sentence: teaching large language models the language of biology. BioRxiv, 2023.2009. 2011.557287 (2024).

10. Baek, S., Song, K. & Lee, I. Single-cell foundation models: bringing artificial intelligence into cell biology. Experimental & Molecular Medicine, 1–13 (2025).

11. Yang, A., et al. Qwen3 technical report. *arXiv preprint arXiv:2505.09388* (2025).

12. Wolf, F.A., Angerer, P. & Theis, F.J. SCANPY: large-scale single-cell gene expression data analysis. Genome biology 19, 15 (2018).

13. Shahapure, K.R. & Nicholas, C. in 2020 IEEE 7th international conference on data science and advanced analytics (DSAA) 747-748 (IEEE, 2020).

14. Korsunsky, I. et al. Fast, sensitive and accurate integration of single-cell data with Harmony. Nature methods 16, 1289–1296 (2019).

15. Nguyen, Q.H. et al. Profiling human breast epithelial cells using single cell RNA sequencing identifies cell diversity. Nature communications 9, 2028 (2018).

16. Wu, S.Z. et al. A single-cell and spatially resolved atlas of human breast cancers. Nature genetics 53, 1334–1347 (2021).

17. Tuong, Z.K. et al. Resolving the immune landscape of human prostate at a single-cell level in health and cancer. Cell reports 37 (2021).

18. Mages, S. et al. TACCO unifies annotation transfer and decomposition of cell identities for single-cell and spatial omics. Nature biotechnology 41, 1465–1473 (2023).

19. Bae, S. et al. CellDART: cell type inference by domain adaptation of single-cell and spatial transcriptomic data. Nucleic acids research 50, e57–e57 (2022).

20. Lin, A. et al. Bridging artificial intelligence and biological sciences: a comprehensive review of large language models in bioinformatics. Briefings in Bioinformatics 26, bbaf357 (2025).

21. Das, A., Kong, W., Sen, R. & Zhou, Y. in Forty-first International Conference on Machine Learning (2024).

22. Li, X. et al. A dynamic single cell-based framework for digital twins to prioritize disease genes and drug targets. Genome medicine 14, 48 (2022).

23. Almási, S., Krenács, T., Krenács, L. & Cserni, G. Galectin-1 expression in breast cancer stroma–prognostic value in triple-negative breast cancer. Pathobiology (2025).

24. Grosset, A.-A. et al. Galectin signatuares contribute to the heterogeneity of breast cancer and provide new prognostic information and therapeutic targets. Oncotarget 7, 18183 (2016).

25. Rajapaksa, U.S., Jin, C. & Dong, T. Malignancy and IFITM3: friend or foe? Frontiers in Oncology 10, 593245 (2020).

26. Yang, M., Gao, H., Chen, P., Jia, J. & Wu, S. Knockdown of interferon-induced transmembrane protein 3 expression suppresses breast cancer cell growth and colony formation and affects the cell cycle. Oncology Reports 30, 171–178 (2013).

27. Gaspard-Boulinc, L.C., Gortana, L., Walter, T., Barillot, E. & Cavalli, F.M. Cell-type deconvolution methods for spatial transcriptomics. Nature Reviews Genetics, 1–19 (2025).

28. Yan, C. et al. Integration tools for scRNA-seq data and spatial transcriptomics sequencing data. Briefings in Functional Genomics 23, 295–302 (2024).

29. Hu, E.J. et al. Lora: Low-rank adaptation of large language models. ICLR 1, 3 (2022).

30. Tang, M. et al. Evaluating single-cell cluster stability using the Jaccard similarity index. Bioinformatics 37, 2212–2214 (2021).

