## Supplementary Figures for "Generative single-cell transcriptomics via large language models"

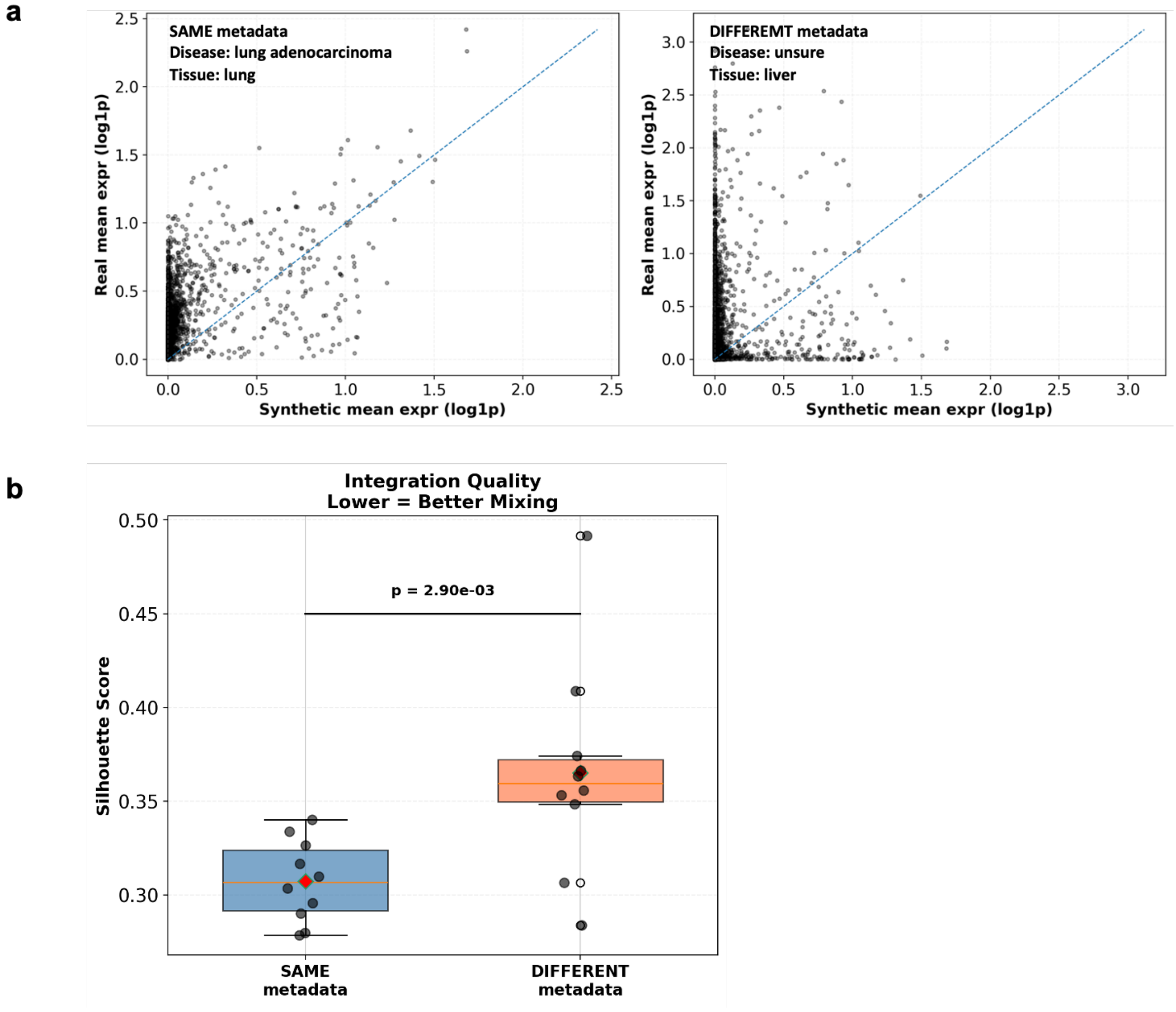


**Supplementary Figure 1 | Additional evaluation of transcriptomic similarity between generated and real scRNA-seq data**

a, Representative examples of gene-wise expression similarity between PGL-generated and real scRNA-seq datasets. Scatter plots show gene-level expression distributions. Comparisons are shown for datasets with matched metadata (same tissue and disease) and mismatched metadata (different tissue and/or disease), illustrating higher concordance in matched conditions.

b, Single-cell–level integration of PGL-generated and real scRNA-seq datasets without batch correction. Silhouette scores quantify the degree of mixing between generated and real cells. Integrations with datasets sharing the same metadata exhibit significantly lower silhouette scores than integrations with mismatched metadata, indicating improved transcriptional similarity.

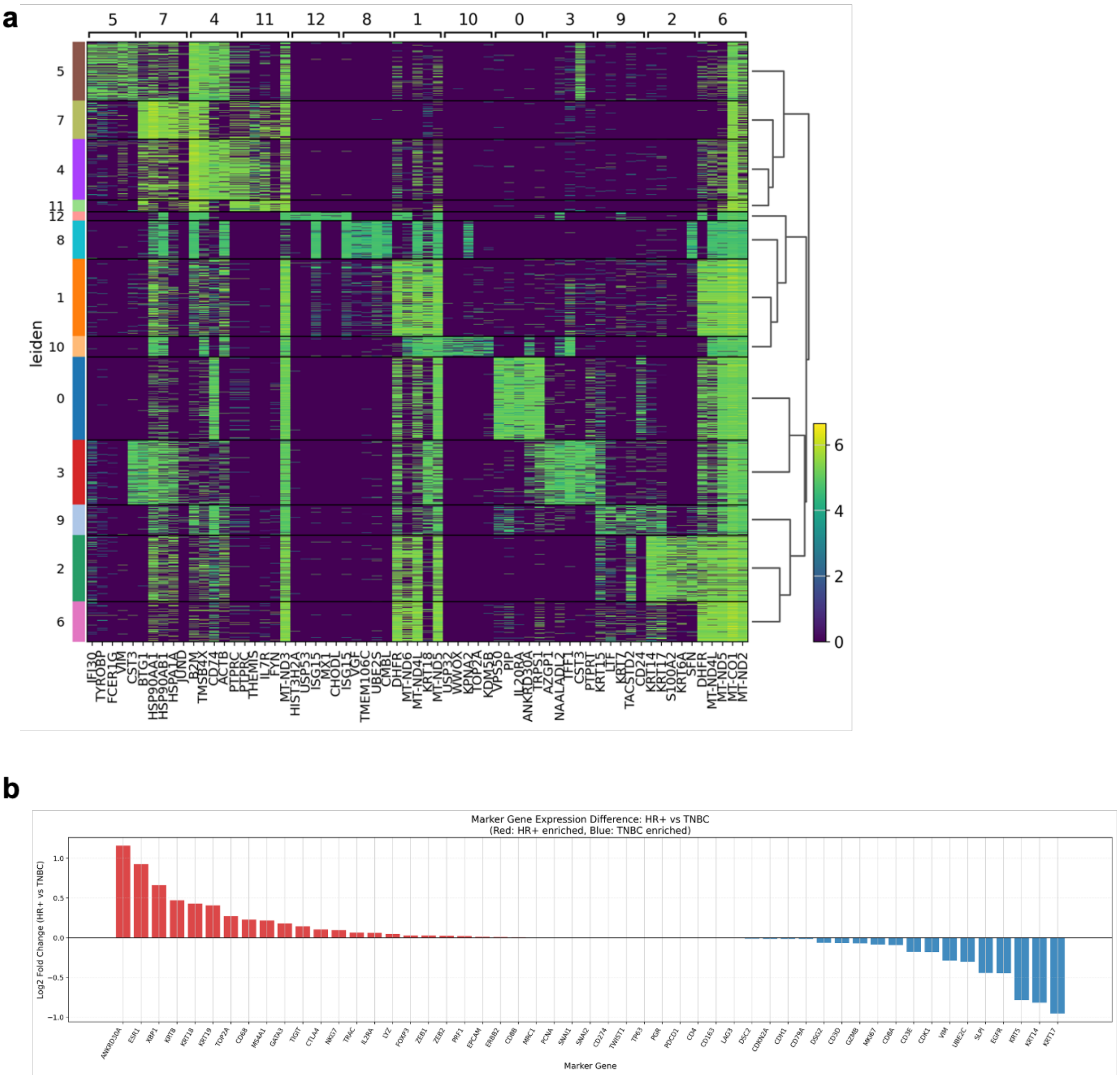


**Supplementary Figure 2 | Additional analyses of breast cancer subtype–specific transcriptomic structure**

a, Heatmap of integrated breast cancer scRNA-seq datasets, colored by clusters and extracted marker gene expression.

b, Canonical gene markers curated luminal and basal gene signatures across generated HR-positive and TNBC datasets. Luminal gene sets are significantly enriched in HR-positive cells, whereas basal gene sets are enriched in TNBC cells, consistent with established breast cancer biology.

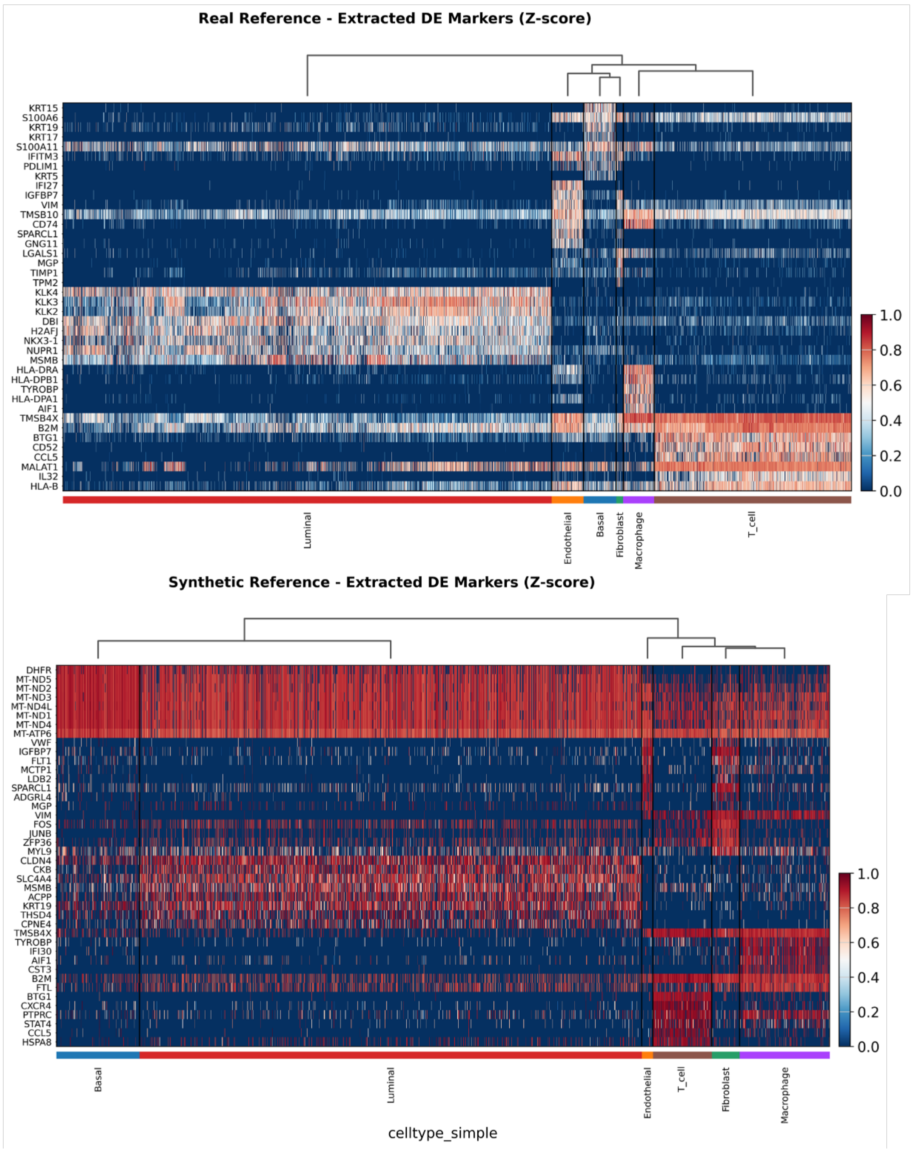


**Supplementary Figure 3 | Marker gene expression used for cell type annotation in prostate cancer datasets**

Heatmaps showing scaled expression of canonical marker genes used to annotate major cell types (epithelial, fibroblast, endothelial, myeloid, and lymphoid) in (upper) PGL-generated prostate cancer scRNA-seq data and (lower) real prostate cancer scRNA-seq atlas data. Marker expression patterns support consistent cell type annotation across synthetic and real datasets.

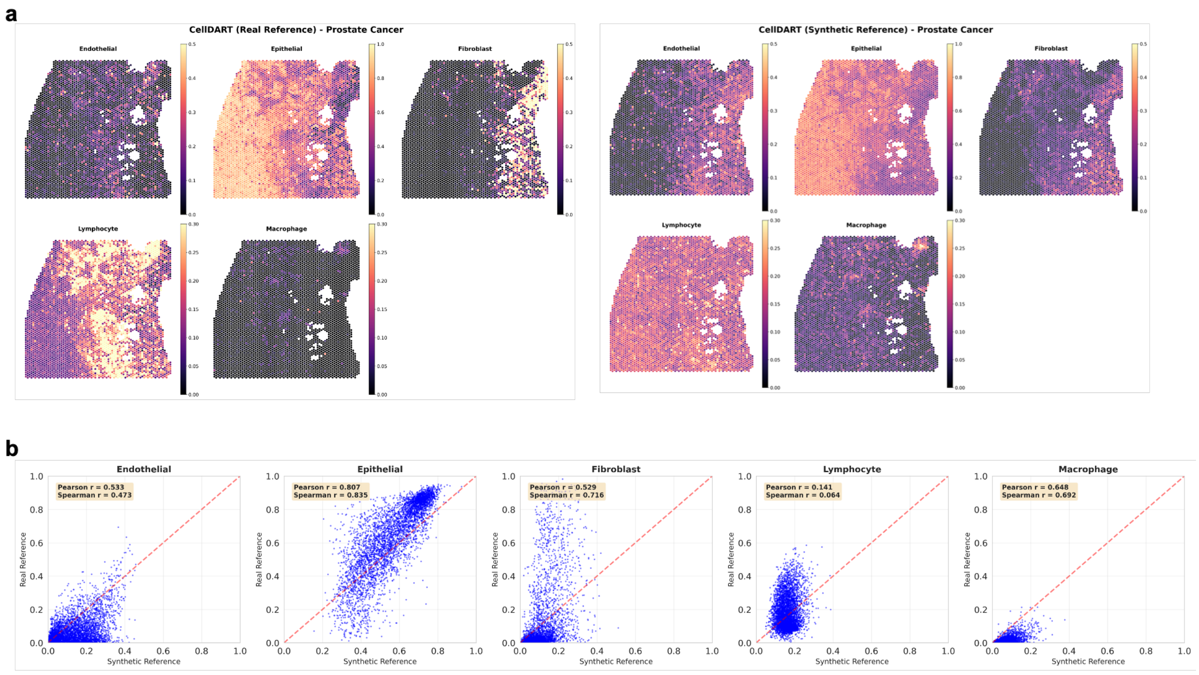


**Supplementary Figure 4 | Spatial cell type mapping using an alternative deconvolution method**

a, Spatial cell type proportion maps inferred using CellDART with PGL-generated scRNA-seq data as reference for prostate cancer Visium samples.

b, Spot-wise correlation analysis comparing cell type proportions inferred using synthetic versus real scRNA-seq reference datasets with CellDART. Significant positive correlations are observed across all major cell types, indicating robustness of synthetic references across deconvolution frameworks.

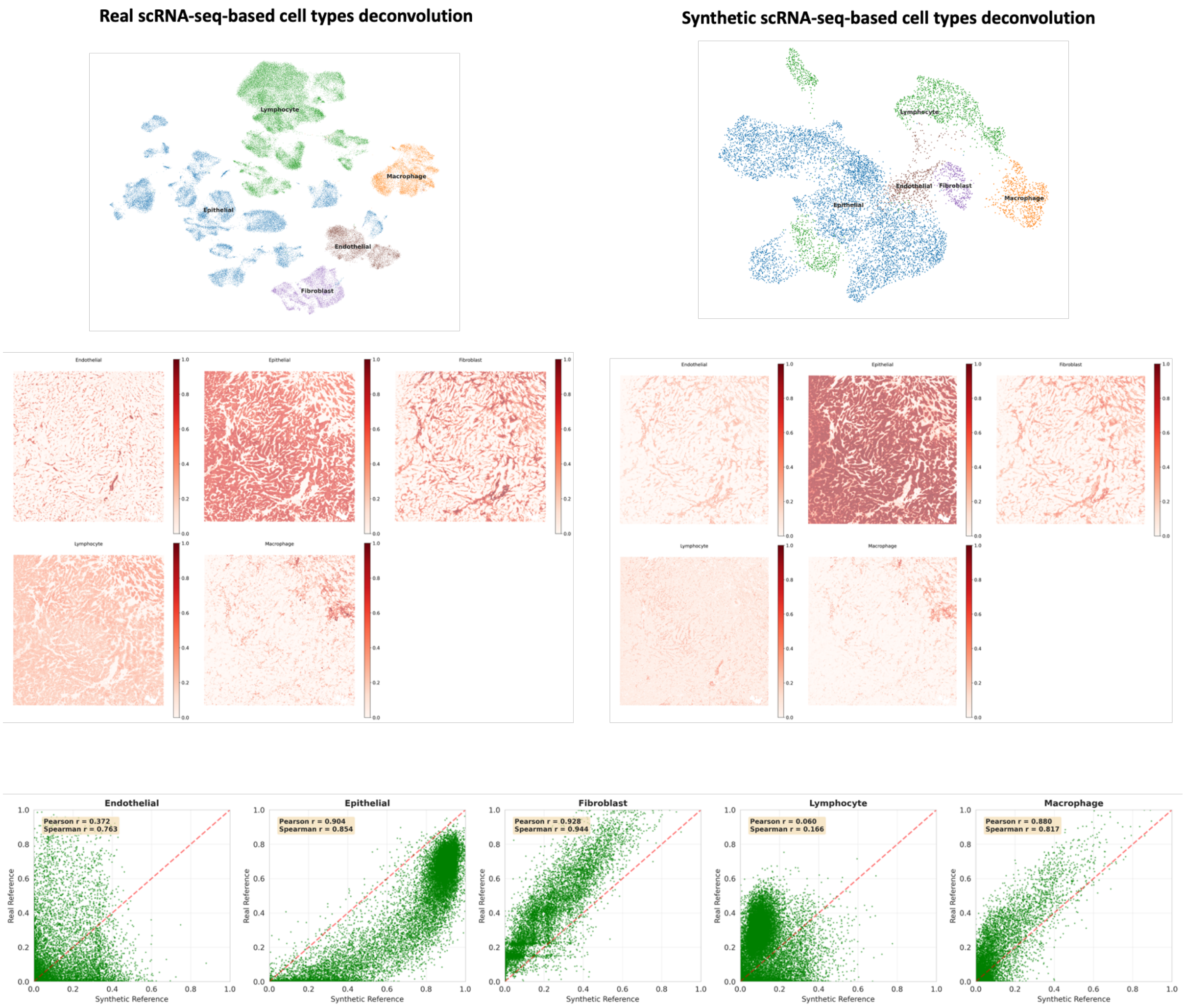


**Supplementary Figure 5 | Spatial mapping performance in high-resolution Visium HD data**

Comparison of spatial cell type mappings obtained using PGL-generated and real scRNA-seq reference datasets in Visium HD breast cancer samples (16-µm spatial bins). Spot-wise correlations of inferred cell type proportions demonstrate strong agreement across reference types, confirming applicability of synthetic references at higher spatial resolution.
